# Aging Is Associated with Reduced Topographic Differentiation Between Speaking and Silent Reading

**DOI:** 10.64898/2026.10.14.756359

**Authors:** Alexis Basciano, Ayoub Daliri, Sara-Ching Chao, Corianne Rogalsky

**Affiliations:** College of Health Solutions, Arizona State University, Tempe, AZ, United States

**Keywords:** Aging, Network Specificity, Topographic Maps, Global Dissimilarity

## Abstract

Aging affects how neural activity is organized and specialized, but its impact on neural differentiation across language tasks remains unclear. In this study, we investigated whether aging influences neural differentiation during speaking and silent reading. Fifteen young adults and 15 older adults completed speaking and silent-reading tasks in a delayed-response paradigm while we collected electroencephalography (EEG) data. We examined EEG signal strength using time-resolved global field power (GFP), spatial configuration using topographic analysis of variance (TANOVA), and task-specific neural differentiation using the global dissimilarity index (DISS). We also examined the relationship between neural and behavioral measures, including digit-span scores and speech onset time. We found that speaking was associated with higher GFP than silent reading in the 388–800 ms interval, with no significant effects of group or group × task interaction, indicating similar EEG strength across groups. TANOVA revealed age-related differences in scalp topography in the 229–334 ms and 377–538 ms intervals, a task effect in the 618–723 ms interval, and a group × task interaction in the 562–636 ms interval. Notably, young adults had greater DISS than older adults during the 508–629 ms and 660–800 ms intervals, reflecting more distinct neural configurations for speaking and reading. We did not find significant correlations between DISS or GFP and behavioral measures, and the two groups performed similarly on behavioral measures. Overall, aging was associated with reduced neural differentiation between speaking and silent reading, especially during later processing stages (e.g., motor command preparation), while preserving overall EEG activity and behavioral performance. These findings may suggest that aging reduces neural network specificity (as previously suggested) and alters spatial organization without necessarily impairing behavior.

## Introduction

Older adults experience widespread structural (Giorgio et al., 2010; Liu et al., 2017) and functional (Geerligs et al., 2015; Sala-Llonch et al., 2015; Sun et al., 2011) changes in the brain, particularly within neural networks that support complex behaviors (Deery et al., 2023). A prominent consequence of these age-related changes is reduced network specificity (Song et al., 2014). Network specificity is the extent to which the brain recruits distinct, task-relevant networks for different cognitive or motor functions. Reduced network specificity occurs when the same brain regions or networks are recruited across multiple tasks, leading to greater overlap in neural activity. For example, older adults tend to engage more generalized and overlapping neural systems across cognitive, sensory, and motor domains (Bernard & Seidler, 2012; Carp et al., 2011; Edde et al., 2021; Lalwani et al., 2019; Park et al., 2004). This pattern is particularly evident in language-related processes, including semantic and phonological processing (Diedrichs et al., 2026; Martins et al., 2014). Reduced network specificity may reflect diminished functional specialization, limiting the selective engagement of task-relevant networks.

Alternatively, overlapping neural recruitment may reflect compensatory mechanisms that help maintain performance despite age-related change. Thus, whether reduced network specificity is adaptive or maladaptive in aging remains unclear (Cabeza, 2002; Park & Reuter-Lorenz, 2009; Simmonite & Polk, 2022). A complete characterization of these changes and their functional relevance is critical for distinguishing typical from abnormal brain aging. Although substantial evidence supports reduced neural specificity in older adults, we know much less about its temporal dynamics. Most studies of this nature use functional Magnetic Resonance Imaging (fMRI). While useful for detecting these network changes, this approach is limited by reduced temporal precision due to the hemodynamic response (Menon & Kim, 1999). Thus, it is difficult to capture neural changes in the brain during rapid and complex cognitive tasks.

Electroencephalography (EEG) is a noninvasive neuroimaging method with high temporal resolution that estimates brain responses to complex stimuli through event-related potentials (ERPs). ERPs reflect stimulus- or event-related changes in scalp-recorded electrical activity generated by populations of neurons (Luck, 2014). Traditional ERP analyses focus on differences in component amplitude and latency across experimental conditions (Luck & Kappenman, 2011). For example, the N1 is a negative-going component that typically peaks around 100 ms after stimulus onset and is associated with early sensory processing. Traditional ERP analysis examines how stimulus characteristics modulate the N1 peak and latency. However, ERP components are strongly influenced by development and aging, resulting in substantial age-related differences in waveform morphology. These differences can make comparisons of individual component amplitude and latency, such as the N1, across age groups challenging. Unlike traditional ERP analyses that compare peak amplitude and latency, time-resolved scalp-wide analyses, such as topographic analysis of variance (TANOVA; Koenig et al., 2011; Habermann et al., 2018), can identify when conditions differ in their spatial patterns of scalp activity (Haartsen et al., 2022; Heald et al., 2022; Jie et al., 2025). Such methods take into account data from all scalp electrodes and can provide information about how the brain is engaged across experimental conditions. TANOVA compares scalp topographies across groups or conditions at each time point after normalizing each map by global field power (GFP), which quantifies the overall strength of the EEG signal across the scalp. This normalization allows TANOVA to focus on differences in the spatial distribution of activity rather than differences in overall amplitude. TANOVA then uses nonparametric statistical tests, typically permutation tests, to determine whether the topographic differences between groups and conditions are greater than would be expected by chance. However, because TANOVA examines the group averages, it does not provide a continuous measure of topographic differentiation for individual participants. A global dissociation index (DISS), as described by Murray et al. (2008), quantifies the dissimilarity between EEG topographic maps across conditions for each participant. DISS normalizes each map by its GFP and then calculates the root-mean-square difference between the normalized voltages across all electrodes. Overall, unlike traditional ERP analyses, TANOVA and DISS do not rely on peak amplitude and latency at specific electrodes; instead, they consider all electrodes and the overall strength of the EEG signal (i.e., GFP). While TANOVA determines whether the topographical maps differ across conditions and groups, DISS quantifies the magnitude of that difference at the participant level. Thus, combining TANOVA and DISS approaches has the potential to uncover the temporal and spatial nature of age-related alterations in neural networks involved in complex tasks.

Word reading is a highly complex task that involves several processes, including visual attention and grapheme recognition, phonological decoding (Seidenberg & McClelland, 1989), and, during overt reading (i.e., speaking), the planning and execution of coordinated speech movements (Guenther, 2006). Speaking and silent reading engage overlapping neural networks involved in visual word recognition and phonological processing. However, speaking additionally recruits speech-motor networks required to plan and produce speech (Brumberg et al., 2016; Dietz et al., 2005; Turkeltaub et al., 2002). Aging affects several processes involved in word reading, including visual attention (Curran et al., 2001; Zhuravleva et al., 2014) and motor planning (Bernard & Seidler, 2012; Reuter et al., 2015). However, few studies have directly compared the neural processes involved in speaking and silent reading in older adults. This comparison is challenging because speech preparation and production occur rapidly and overlap in time. A delayed-speaking paradigm addresses this challenge by asking participants to prepare to read a target word aloud and then begin speaking only after a cue. This design separates speech preparation from speech execution and has been widely used to study speech planning (Daliri et al., 2015, 2016, 2018; McGuffin et al., 2020; Merrikhi et al., 2018). When combined with EEG, this paradigm can reveal when neural networks engage during speaking and reading. Importantly, comparing these temporal patterns may provide a sensitive means of determining whether older adults show reduced differentiation between the neural networks engaged in these two tasks.

The primary objective of this study was to determine whether aging reduces neural differentiation during speaking and silent reading. We used a delayed-speaking task in which a word appeared on the screen, and participants prepared to read it either silently or aloud after a visual “go” cue. We recruited young and older adults to examine age-related differences in the temporal dynamics of neural differentiation. We used TANOVA to identify differences in scalp topographic patterns between the speaking and reading tasks during the preparation phase. We also calculated DISS values for the topographic maps of the two tasks at each time point for each participant. We hypothesized that young adults would show greater topographic differentiation between speaking and reading than older adults, particularly during the speech preparation stage. Additionally, we hypothesized that if reduced neural differentiation (especially in older adults) is associated with less efficient cognitive processing, then participants with reduced neural differentiation, as estimated by DISS values, would show poorer cognitive performance.

## Methods

### Participants

We recruited 30 participants, comprising 15 young adults (5 males; 25.15 ± 4.31 years, 18–32 years) and 15 older adults (8 males; 70.46 ± 9.95 years, 49–85 years). All participants demonstrated normal hearing thresholds (<25 dB HL across 250–4000 Hz, ASHA, 1997). All participants reported no preexisting or current neurological, psychological, or speech-language disorders. The Mini-Mental Status Exam (MMSE) (Folstein et al., 1975) was administered to all older adult participants to ensure they did not have preexisting mild cognitive impairment (all scores were greater than 24). All participants (young and older) completed the verbal working memory subsection of the Wechsler Adult Intelligence Scale (WAIS) IV (Wechsler, 2008), including forward digit span, backward span, and sequencing. Experimental and recruitment procedures were preapproved by Arizona State University’s (ASU) Institutional Review Board.

### Apparatus

Data collection for the entire session occurred in a sound-attenuated booth. Participants were seated in front of an Alienware computer monitor (24 inches, 240 Hz refresh rate), while wearing insert earphones (Etymotic Research ER2). The computer monitor was controlled by a computer outside the sound booth, which presented task instructions and target words throughout the experiments using Psychtoolbox in MATLAB (Kleiner et al., 2007). A microphone (Shure SM58) was placed 15 cm from the participant’s mouth, angled at 45 degrees. During speaking blocks, the speech signal from the microphone was amplified via an ART TubeOpto 8 and sent to a MOTO interface (8pre USB). Participant speech was then amplified (via a Samson S-phone) to be 5 dB louder than the original signal (Chao et al., 2019) and played back to participants binaurally through the insert earphones.

Electroencephalography (EEG) signals were recorded continuously throughout the experimental session using a BrainVision 64-electrode Ag/AgCl setup with a 10-10 system (Oostenveld & Praamstra, 2001). At the start of each session, participants were fitted with an appropriately sized 64-electrode actiCAP (with SNAP holders), and the cap was properly positioned on the scalp. Passive Ag/AgCl electrodes applied to the face were used to monitor eye blinks and muscle activity. We recorded electrooculogram (EOG) using two electrodes: one positioned at the right supraorbital area and the other at the left infraorbital area. Additionally, we recorded electromyography (EMG) using two passive electrodes placed over the left and right orbicularis oris muscles. EEG data were recorded continuously at a 1000 Hz sampling rate throughout the session using PyCorder software on a separate computer. Synchronously with EEG, the speech signal from a second microphone (Shure MVL Lavalier) was recorded.

### Procedure

Cognitive assessments were conducted at the beginning and midpoint of the experimental session to reduce cognitive fatigue. At the start of the session, a trained experimenter administered the Mini-Mental-Status Exam (MMSE) to all older adult participants. At the halfway point of data collection, the experimenter entered the sound booth and administered the Digit Span subtest of the WAIS-IV Working Memory Index (WMI) (Wechsler, 2008), which includes the Digit Span Forward, Digit Span Backward, and Digit Span Sequencing. In the Forward Span, participants repeated numbers in the order they were presented, as a measure of attention and short-term memory. In the Backward Span, participants repeated the numbers in reverse order, a measure of working memory. In the Sequencing subtest, participants repeated numbers in ascending order, a measure of working memory capacity and the ability to mentally reorganize and manipulate information. During this time, participants were instructed to remove their earphones to ensure they heard the WAIS test items. Afterward, the experimenter assisted participants in repositioning the headphones to continue collecting neural data. Following the cognitive assessment, participants completed several blocks of different experimental tasks in random order.

### Experimental tasks: Speaking and reading

First, participants completed a practice task to become familiar with the overall experiment. The task structure throughout the experiment was a delayed-response paradigm in which participants either said a word aloud or read it silently after a cue. Across all paradigms, target words were monosyllabic and followed a consonant-vowel-consonant structure with the vowel /ɛ/ (e.g., “head”). Participants first completed a 2-minute practice to become accustomed to the paradigm. At the start of each trial, the target word appeared in the center of the screen (capitalized in red on a gray background). After 800 ms, the word turned green, signaling participants to say it aloud. The green word remained on the screen for 2000 ms. Afterward, visual feedback on speech production was presented for 750 ms. The feedback screen indicated whether speech onset, intensity, and duration were out of range. The feedback screen was followed by a gray screen between trials, with jittered inter-trial intervals of 1 to 1.5 s. Subsequent tasks did not include these feedback screens between trials. In the subsequent tasks, participants completed blocks of speaking and reading. Each block consisted exclusively of either speaking or reading trials. Task procedures were identical to the practice: participants read words following a “go” signal (word changing from red to green), except that in reading blocks they silently read the word without any movement. Each block of the experimental task took about 6 minutes, and the order of the experimental tasks was randomized. During the experimental session, we also collected additional speaking blocks as part of a separate study.

### Data Analysis

The EEGLAB toolbox was used to process EEG signals. We began by merging all blocks into a single dataset per participant for each condition, one for speaking and one for reading. Each subject’s aggregated dataset was then filtered using a zero-phase Hamming-window FIR filter (high-pass cutoff: 0.5 Hz; low-pass cutoff: 50 Hz). Next, we cleaned the data and rejected artifacts. In this stage, EEGLAB selectively removed bad channels, including those with a correlation below 0.85 with neighboring channels after reconstruction, and noisy channels that exceeded 4 standard deviations from the mean signal. Next, we used Artifact Subspace Reconstruction (ASR), implemented in EEGLAB (Mullen et al., 2015), to automatically remove local, transient, high-amplitude deviations in the continuous data while preserving the underlying neural signal. For time windows with high variance, the function reconstructed the signal using the remaining, cleaner channels. To simplify subsequent group-level comparison, we employed spherical spline interpolation (Perrin et al., 1989) to reconstruct previously dropped noisy channels. Following filtering and cleaning, each participant had one dataset for speaking and one for reading.

We then applied Independent Component Analysis (ICA) to the continuous EEG signal to identify independent components associated with artifacts, including eye blinks and muscle activity. For this purpose, we used EEGLAB to categorize each independent component. Components with over a 90% likelihood of being related to eye movement, muscle activity, or channel noise were identified and removed from the data to ensure the neural signal remained clear. The dataset was then average-referenced, as this is essential for calculating TANOVA and DISS analyses. The continuous reading and speaking blocks were then epoched, starting 200 ms before the red word appeared and ending 200 ms after the green go signal appeared. We then corrected for baseline activity by removing the average amplitude from the first 100 ms of the entire 1200 ms epoch. These aggregated, baseline-corrected signals for each condition (speaking and reading) for each participant were used in subsequent ERP, topographical, and GFP analyses.

### Topographic Analysis

Multichannel analyses of the processed EEG data were conducted using the RAGU (Randomization Graphical User Interface) software (Koenig et al., 2011; Habermann et al., 2018). Initially, a topographic consistency test verified consistent activation throughout the 1200 ms epoch within each group and task. RAGU’s TANOVA assesses whether differences across experimental conditions are present in the scalp electric field’s spatial distribution, independent of overall field strength variations. For each time point, scalp potential maps are normalized by their global field power (GFP) to remove amplitude differences while preserving relative spatial activity across electrodes. The dissimilarity between normalized maps across tasks and groups is then measured using a global map-dissimilarity metric. Significance is determined using permutation tests, in which task labels are shuffled within each participant to create a null distribution of the topographic difference statistic at each time point. Comparing observed statistics to this distribution yields a time-resolved significance estimate. Therefore, TANOVA detects periods when tasks differ in topographic configuration. RAGU applies a similar permutation approach to assess the main effect of group and the group-by-task interaction at each time point. To address multiple comparisons across the time series, RAGU calculates global permutation-based statistics that evaluate the overall significance pattern over time, such as the Global Duration metric, which assesses whether the extent or strength of effects exceeds chance expectations. These global statistics help control the multiple-testing problem while identifying topographic differences. Overall, TANOVA identifies time intervals during which the spatial configuration of the scalp topographic maps differs significantly between groups, tasks, or their group-by-task interaction.

While TANOVA detects group-level significance, it doesn’t quantify individual topographical differences. We therefore calculated a global dissimilarity (DISS) score (Murray et al., 2008) by comparing each participant’s speaking and reading maps. First, data were GFP-normalized to remove signal-strength effects. Squared electrode differences were averaged, and the square root of these averages yielded the DISS value, reflecting topographic similarity between conditions for each participant. DISS scores range from 0 (identical maps) to 2 (inverse maps). To explore age-related differences, DISS values were compared between young and older adults at each time point using a nonparametric permutation test. Group differences in DISS were calculated, labels were shuffled randomly, and differences were recalculated to form a null distribution. Permutation p-values were derived by comparing observed differences to this null. To control for multiple time points, a maximum cluster-based permutation test (Maris & Oostenveld, 2007) was used: group labels were shuffled, continuous clusters exceeding an uncorrected threshold of .05 were identified, and the test statistic sums from these clusters were used to build an empirical null distribution. Clusters exceeding the 95th percentile of this distribution were considered significant. Lastly, to compare overall EEG signal strength across tasks and groups, we examined differences in GFP using nonparametric permutation tests at each time point and applied the same max-cluster method to identify significant intervals.

## Results

### Examining global field power

We initially compared GFPs across tasks and groups. GFP reflects the overall EEG signal strength across all scalp electrodes. Because we used GFP to normalize EEG signals before analyzing topographical differences, it’s essential to assess GFP variation across tasks and groups. Figures 1A and 1B show the GFP waveforms for the speaking and reading tasks in young and older adults, respectively. Our nonparametric tests showed a significant main effect of task from 388 to 800 ms, with larger GFPs during the speaking task than during the reading task (p < .001). However, neither the main effect of age group (p > .157) nor the group-by-task interaction (p > .0936) was statistically significant. Figure 1C presents data for the significant interval (388–800 ms) for both tasks and groups.

**Figure 1.**
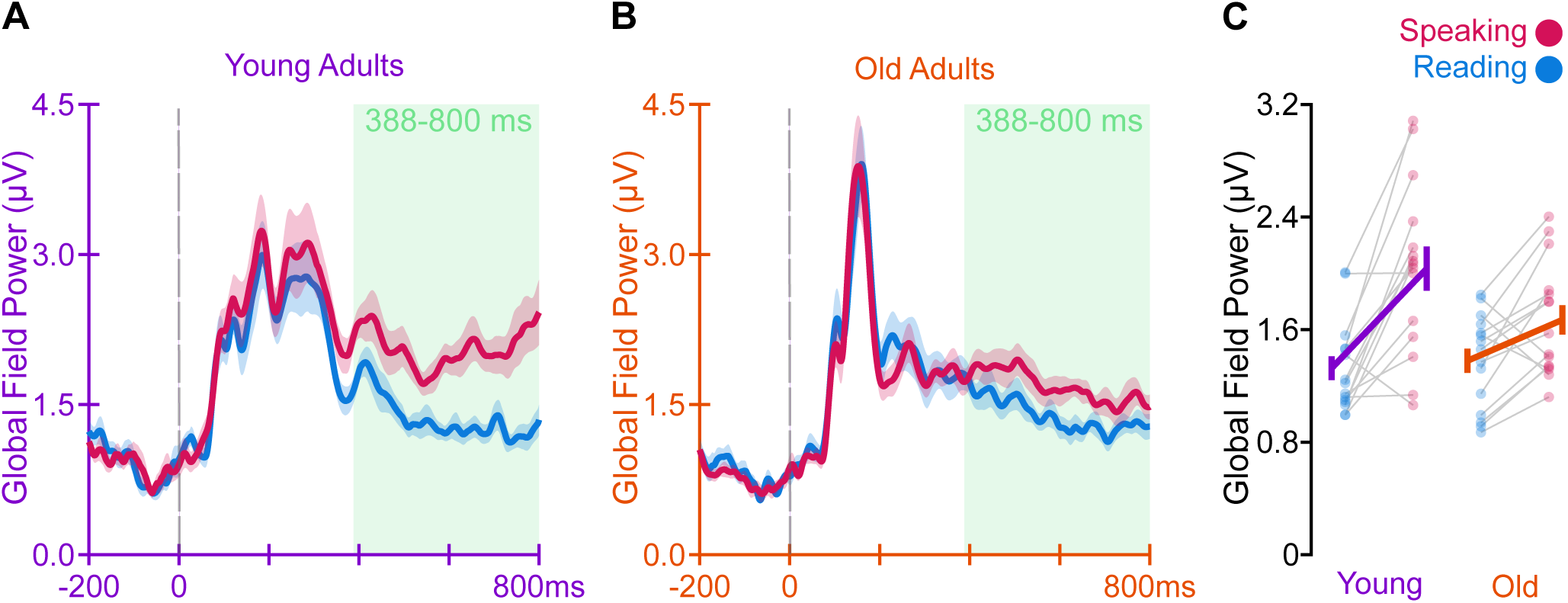
Global field power (GFP) during speaking and silent reading in young and older adults. GFP reflects the overall strength of the EEG signal across all scalp electrodes. (A) Grand-average GFP time courses for the speaking and reading conditions in young adults. (B) Grand-average GFP time courses for the speaking and reading conditions in older adults. Shaded areas indicate the standard error of the mean. Nonparametric statistical analysis revealed a significant main effect of task in the 388–800 ms window, as indicated by the green box. (C) GFP values were averaged across the significant 388–800 ms interval for each participant and experimental condition. Speaking elicited significantly greater GFP than silent reading, whereas GFP did not differ significantly between young and older adults, and there was no significant age × task interaction. Error bars indicate the standard error of the mean. The gray dashed line indicates the onset of word presentation (0 ms).

### Examining topographical configurations

We performed TANOVA to evaluate topographic differences across the experimental tasks. TANOVA detects time periods when the spatial configurations of scalp topographic maps differ significantly across groups, tasks, or their interaction. Our results revealed a significant main effect of group at two intervals: 229–334 ms (p = .008) and 377–538 ms (p = .001). Additionally, a significant main effect of task was observed from 618–723 ms (p = .014), along with a group-by-task interaction at 562–636 ms (p = .009). Figure 2 displays the average topographic maps for each significant effect across these intervals. While TANOVA identifies differences in EEG spatial distribution, it does not quantify the magnitude of those differences. To address this, we computed DISS values to compare dissimilarities between topographic maps during reading and speaking tasks for each group. Figure 3 shows the DISS waveform throughout the epoch. Nonparametric analysis revealed that the maps of young adults were more dissimilar than those of older adults in two intervals: 508–629 ms (p = .004) and 660–800 ms (p < .001). Figure 3B presents the average DISS values for each participant and the group averages in these intervals.

**Figure 2.**
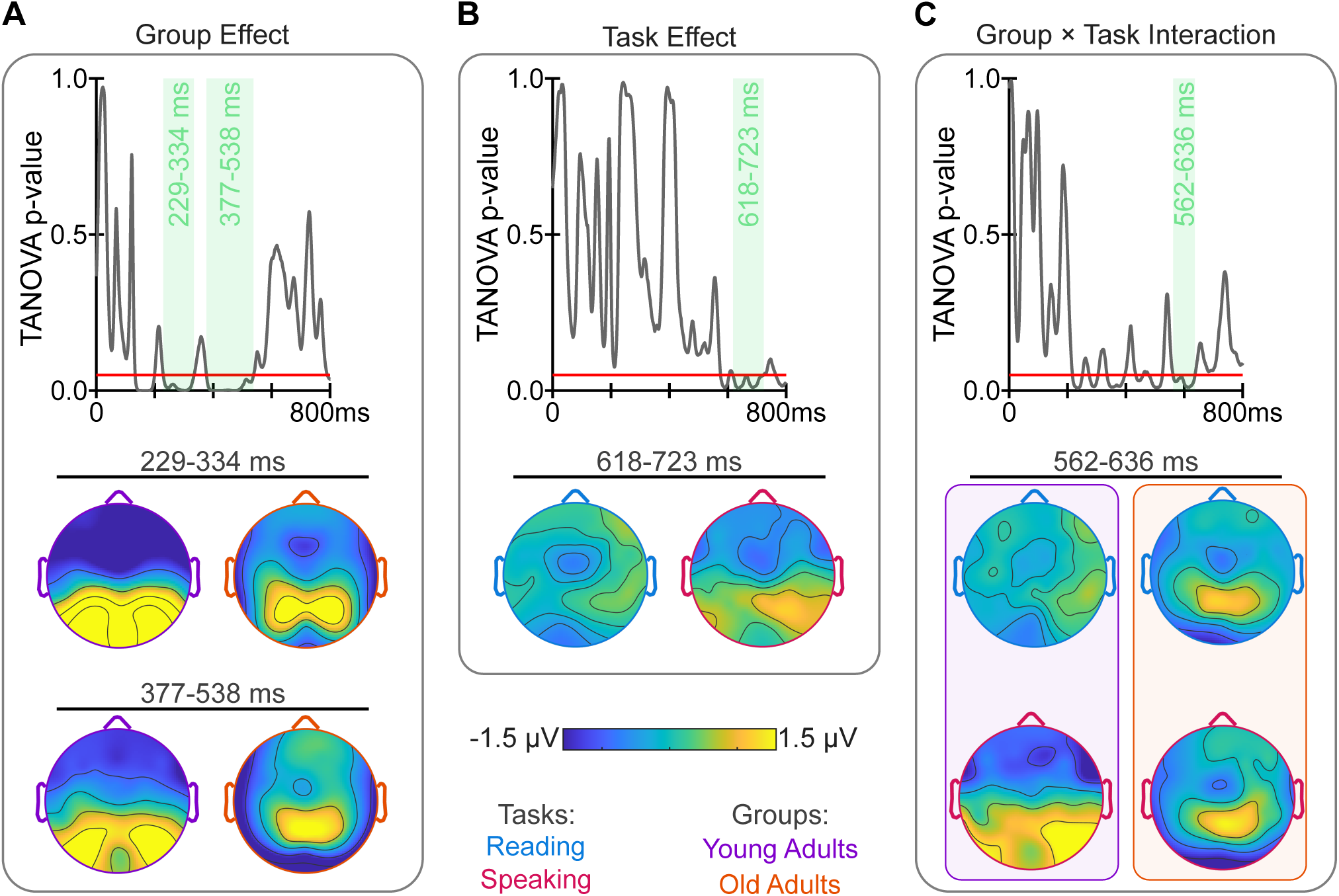
Topographic analysis of variance (TANOVA) of EEG scalp topographies during speaking and silent reading in young and older adults. TANOVA was used to identify time intervals during which the spatial configuration of the scalp electric field differed significantly by age group, task, or their interaction. (A) Group effect. The TANOVA p-value time course shows the statistical significance of the age-group effect across the analysis epoch. Two significant age-related intervals were identified at 229–334 ms and 377–538 ms, as highlighted by the green boxes. Average scalp topographies are shown separately for young and older adults for each significant interval. (B) Task effect. The TANOVA p-value time course indicates a significant difference in scalp topographic configuration between speaking and silent reading from 618–723 ms. Average scalp topographies for the speaking and reading conditions are shown for this significant interval. (C) Group × task interaction. TANOVA revealed a significant age group × task interaction from 562–636 ms, indicating that the effect of task on scalp topographic configuration differed between young and older adults. Average topographies are shown for young and older adults in the speaking and reading conditions during this interval. Color bars represent the scalp voltage distributions. The color of the map boundary (rim) denotes the corresponding experimental task and age group. The red line in the TANOVA p-value panels indicates the significance threshold of p = .05.

**Figure 3.**
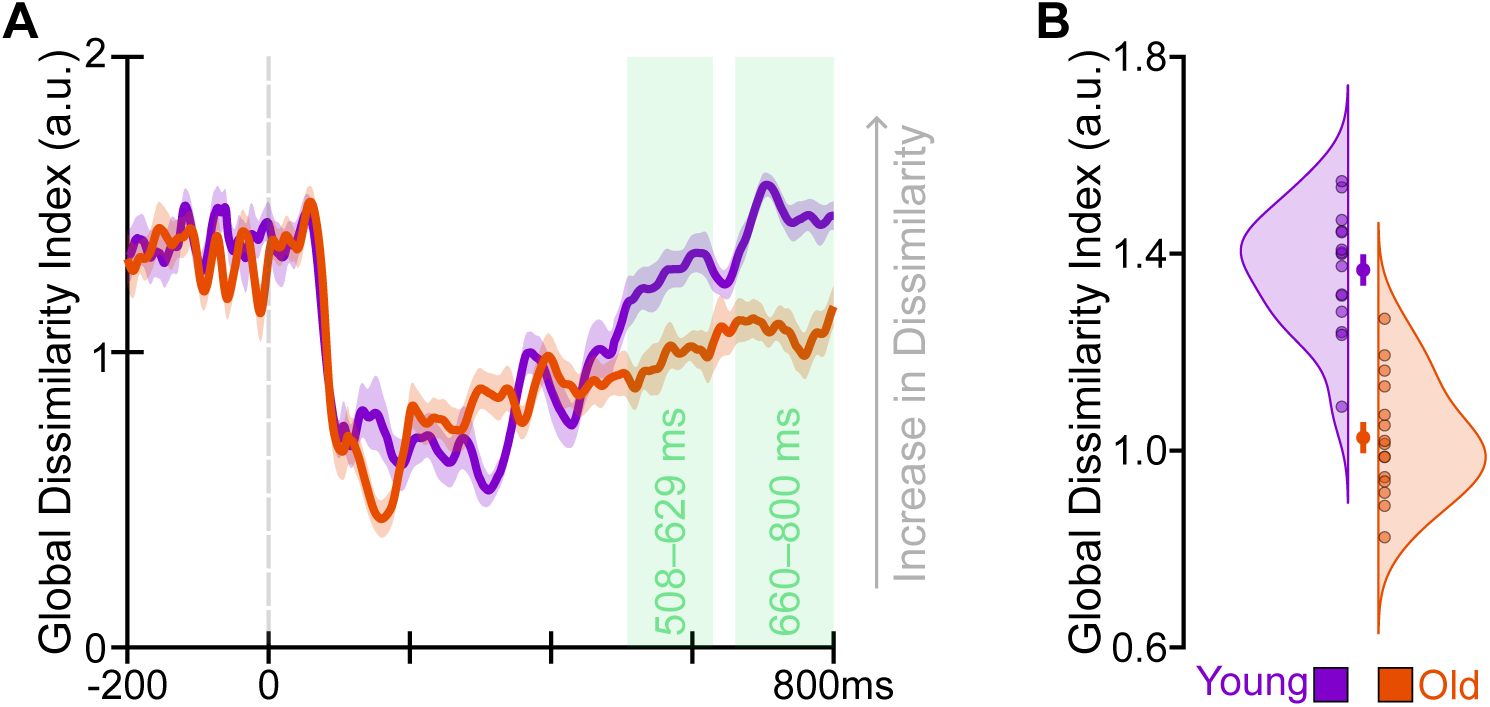
Global dissimilarity (DISS) between speaking-and reading-related EEG topographies in young and older adults. DISS quantifies the dissimilarity between GFP-normalized scalp topographic maps for the speaking and silent reading conditions within each participant, with higher values indicating greater topographic dissimilarity between the two task conditions. (A) Time-resolved DISS values across the analysis epoch for young and older adults. Green-shaded intervals indicate periods when DISS differed significantly between age groups. Nonparametric statistical analysis revealed significantly greater DISS in young adults than in older adults in the 508–629 ms and 660–800 ms intervals, indicating greater speaking–reading topographic differentiation in young adults during these intervals. (B) DISS values averaged within each significant interval for individual participants and their corresponding group-level distributions. Error bars indicate the standard error of the mean. The gray dashed line indicates the onset of word presentation (0 ms).

### Correlational analysis

We conducted several correlational analyses to examine potential associations among GFP, DISS, and behavioral measures, including digit span scores and speech onset times. None of these analyses showed statistically significant relationships. We also examined group differences in these behavioral measures, but again, no significant differences were found.

## Discussion

This study aimed to assess whether aging impacts neural differentiation during speaking and silent reading. Using a delayed-speaking paradigm and time-resolved EEG topographic analyses, we compared overall neural activity strength and spatial patterns between young and older adults during preparation for both tasks. Our nonparametric analyses identified three key findings. First, speaking elicited higher GFPs than silent reading, ranging from 388 to 800 ms, indicating greater EEG signal strength during speaking, with no age-group differences. Second, topographic maps revealed age-related differences at 229–334 ms and 377–538 ms, task-related differences at 618–723 ms, and an age-by-task interaction at 562–636 ms. Third, DISS values were higher in young adults than in older adults in the 508–629 ms and 660–800 ms windows, indicating more differentiated neural configurations in the young. These measures did not correlate with behavioral data. Overall, the results suggest that aging reduces neural differentiation between speaking and silent reading, especially during later processing stages.

Our first finding was that speaking was associated with greater overall scalp electrical field strength than silent reading between 388 and 800 ms. GFP provides a measure of the overall strength of the EEG electrical field across electrodes but does not characterize its spatial configuration or identify the underlying neural sources. Therefore, higher GFP cannot, by itself, distinguish between stronger activity within the same neural populations and the recruitment of additional or different neural populations. In the present study, the greater GFP observed during speaking may be consistent with the greater processing demands of the speaking condition, which involves visual word recognition, phonological processing, and preparation for speech production. In particular, the increased GFP may reflect greater overall neural engagement associated with preparing speech motor commands. Importantly, there was no significant main effect of age group or age group × task interaction in GFP. Thus, older adults did not show a general reduction in overall EEG field strength, nor did the speaking-related increase in GFP differ significantly between age groups. This finding is particularly important for interpreting subsequent topographic analyses, as it indicates that the age-related differences observed in scalp topography cannot be attributed solely to weaker overall EEG responses in older adults. Rather, the results suggest that the age-related differences observed in later analyses are more closely related to differences in the spatial organization and task-related differentiation of neural activity than to differences in overall field strength. Overall, the GFP results show a robust difference in the magnitude of EEG signals between speaking and silent reading, consistent across age groups.

Our second main finding was that both age and task influenced the spatial configuration of the EEG electric field across electrodes. TANOVA revealed a significant main effect of task between 618–723 ms, indicating that speaking and silent reading produced distinct scalp topographic patterns during this period. Because TANOVA was performed on GFP-normalized topographies, this effect reflects changes in the spatial arrangement of the scalp electric field rather than overall EEG signal strength. These distinct topographies likely reflect different neural processes involved in each task. While both tasks involve processing the visual word, speaking additionally requires preparing for speech, including motor commands. This late topographic difference coincides with task-specific neural activity during speech preparation. However, scalp-level TANOVA cannot identify the exact neural sources; thus, topographic differences cannot directly reveal which neural populations or networks are involved. TANOVA also identified significant age effects between 229–334 ms and 377–538 ms, with young and older adults showing different topographies during these periods. Visual maps suggest that young adults have a broader electrical field, whereas older adults show more localized patterns. These visual differences are descriptive rather than definitive evidence of neural localization, but they indicate age-related differences in the organization of task-related neural activity. Notably, the two age-related effects occurred at different times, suggesting that age influences neural topography across multiple processing stages rather than continuously over the 229–538 ms interval. Finally, a significant age group × task interaction between 562–636 ms showed that the effect of task on topography differed between groups. As shown in Figure 3C, young adults exhibited more distinct speaking and reading topographies than older adults during this period. This result is consistent with the DISS analysis, which directly showed greater topographic dissimilarity between speaking and reading in young adults during the 508–629 ms and 660–800 ms intervals. Overall, these findings suggest that aging reduces neural differentiation between speaking and silent reading, consistent with broader evidence of decreased neural network specificity with age. Previous research indicates that older adults tend to activate more overlapping or generalized neural systems across cognitive, sensory, and motor tasks (Bernard & Seidler, 2012; Edde et al., 2021; Lalwani et al., 2019; Park et al., 2004). Similar age-related neural changes have been observed in language processing tasks involving semantics and phonology (Diedrichs et al., 2026; Martins et al., 2014). Our results extend this understanding by showing age differences in the spatial organization and task-related neural differentiation with high temporal resolution during speaking and silent reading.

Our third key finding, which strongly supports the main hypothesis, was that young adults showed significantly higher DISS than older adults during the 508–629 ms and 660–800 ms intervals. DISS assesses how differently the GFP-normalized topographic maps for speaking and reading appear within each participant, with higher values indicating greater differences in neural topography. The higher DISS in young adults indicates more distinct neural patterns for these tasks, whereas the lower DISS in older adults suggests more similar neural configurations. Along with age-related differences in TANOVA topographies, this supports the idea that neural network specificity decreases with age, as evidenced by less differentiation between speaking and silent reading. This interpretation is consistent with previous reports linking aging to reduced functional specialization and greater overlap in neural systems across cognitive, sensory, and motor functions (Bernard & Seidler, 2012; Carp et al., 2011; Edde et al., 2021; Lalwani et al., 2019; Park et al., 2004; Song et al., 2014). Importantly, our results add temporal detail, showing that the age difference in neural differentiation was specific to the 508–629 ms and 660–800 ms windows, indicating that reduced differentiation occurs at particular time points rather than as a constant overall difference. The later timing might relate to speech preparation stages; however, our methods cannot specify the exact processes involved. The most cautious conclusion is that age differences in neural differentiation between speaking and reading develop during the later stages of the neural response.

While our findings point to reduced neural differentiation in older adults, an important question is what functional consequences this reduction may have. To address this question, we examined relationships between the neural measures (DISS and GFP) and behavioral measures (digit-span scores and speech onset time). However, we found no significant relationships between the neural and behavioral measures, and there were no significant age-group differences in the behavioral measures. These findings impose an important constraint on interpretations of reduced neural differentiation, as they do not support a simple association between reduced topographic differentiation and poorer behavioral performance. One possibility is that reduced neural differentiation reflects an age-related change in neural organization that does not necessarily compromise behavioral performance. Older adults may maintain comparable performance despite less distinct neural configurations for speaking and reading, suggesting that changes in neural specificity can occur independently of observable behavioral decline. Another possibility is that older adults achieve comparable performance through alternative patterns of neural organization, such that reduced differentiation reflects a shift in the organization of neural resources rather than a loss of functional capacity. It is also possible that the behavioral measures used in the present study were not sufficiently sensitive to the processes reflected by the measures of neural differentiation. Digit-span scores provide an index of working memory capacity, whereas speech onset time reflects response initiation, and neither measure directly captures the processes underlying the differentiation between speaking and silent reading. Thus, the absence of a neural-behavioral relationship does not necessarily suggest that reduced neural differentiation has no functional consequences. Finally, the preservation of behavioral performance despite reduced neural differentiation could be consistent with adaptive or compensatory recruitment of overlapping neural resources. Such an interpretation is consistent with models proposing that older adults may recruit additional or overlapping neural resources to maintain performance despite age-related changes in neural organization (Cabeza, 2002; Park & Reuter-Lorenz, 2009; Simmonite & Polk, 2022). However, the present findings do not provide direct evidence for compensation, and it would therefore be premature to conclude that the reduced differentiation observed in older adults is compensatory. Future studies using behavioral measures more closely aligned with language processing and speech preparation may help clarify the functional significance of age-related reductions in neural differentiation.

Several limitations should be taken into account when interpreting these results. First, the study involved only 15 young and 15 older adults. While significant age-related differences were observed in the topographic measures, larger samples across a broader age range are necessary to confirm the robustness and generalizability of these findings and to increase statistical power to detect neural–behavioral links. Second, the current analysis describes differences in electrical topography at the scalp level rather than pinpointing the exact neural sources. Methods like TANOVA and DISS reveal the spatial organization and differentiation of neural activity, but they cannot specify which brain regions or networks are involved. Future studies combining EEG with source localization or other neuroimaging techniques may help identify the neural generators responsible for these age-related differences. Third, the stimuli in the speaking and reading tasks were monosyllabic words. Their simplicity likely limited cognitive and linguistic demands, possibly reducing performance variability and decreasing sensitivity to neural-behavioral relationships. Using more complex stimuli or tasks with higher cognitive load could better test the functional significance of age-related neural differences. Lastly, all older participants scored above 24 on the MMSE, indicating cognitively healthy aging. Therefore, these findings primarily reflect neural changes associated with healthy aging and may not apply to individuals with mild cognitive impairment or neurodegenerative conditions. Future research comparing healthy aging with pathological aging could clarify whether the extent and timing of reductions in neural differentiation change progressively as cognitive decline progresses.

In conclusion, the present study examined age-related differences in neural differentiation during speaking and silent reading using time-resolved EEG topographic analyses. Our findings showed that aging is associated with reduced neural differentiation between speaking and silent reading, despite comparable overall EEG signal strength and behavioral performance. Although speaking elicited greater GFP across age groups, TANOVA revealed age-related differences in the spatial organization of neural activity, and DISS showed less distinct speaking- and reading-related topographies in older adults during later processing stages. These findings are consistent with reduced neural network specificity with aging and suggest that age-related changes in neural organization may occur without evident behavioral decline.

## Acknowledgments

This work was supported by a grant from the National Institutes of Health awarded to A. Daliri (R01-DC019905 and R01-DC020162). The content is solely the responsibility of the authors and does not necessarily represent the official views of the National Institutes of Health.

## Data Availability Statement

The dataset collected for the current study is available upon reasonable request from the author (A. Daliri).

